# Identification of 4-Amino-2-Nitrophenol as a novel inducer of phenotypic antibiotic resistance in *Escherichia coli*: roles of Lon protease and its substrate MarA

**DOI:** 10.1101/2025.02.26.640252

**Authors:** Santhi Sanil Nandini, Sirisha Jagdish, Subinoy Rana, Dipankar Nandi

## Abstract

In prokaryotes, the energy-dependent protein degradation is controlled primarily by two ATP-dependent proteases, Lon and Clp. This study investigates the roles of the Lon protease in the metabolism of 2,4-dinitrophenol (2,4-DNP), a toxic industrial compound, in *Escherichia coli* (*E. coli*). During the study, an observation was made that the absence of Lon protease resulted in an enhanced conversion of yellow coloured 2,4-DNP to a reddish-brown product. This study aims to characterise the compound observed in the media with wild type (WT) and Δ*lon* strains, understand the mechanisms of 2,4-DNP conversion and decipher the roles of Lon protease in the conversion of 2,4-DNP. UV-visible and LC-MS analyses revealed differences in the conversion products between the WT and Δ*lon* strains. One of the substrates of Lon protease is MarA, a transcription factor. Growth studies with different mutants and trans-complemented strains demonstrated MarA-dependent conversion. The bathochromic shift of spectral peaks suggested a reduction process and possible involvement of nitroreductase enzymes. Indeed, the expression of two nitroreductases, *nfsA* and *nfsB*, increased with 2,4-DNP and was dependent on MarA. Importantly, the production of the reddish-brown product was lower in strains lacking *nfsA* or *nfsB*. Finally, LC-MS analysis identified one of the conversion products of 2,4-DNP to be 4-Amino-2-nitrophenol (4,2-ANP). Dose studies with purified 4,2-ANP demonstrated that it did not lower the growth of *E. coli* (unlike 2,4-DNP) but induced phenotypic antibiotic resistance (like 2,4-DNP). This study contributes to our understanding of biological treatment of nitroaromatics and may offer insights into environmental pollution mitigation strategies.

**Importance:** This study identifies the roles of Lon protease and its substrate MarA in inducing nitroreductases, NfsA and NfsB, in reducing toxic 2,4-DNP to less toxic 4,2-ANP, a novel inducer of phenotypic antibiotic resistance. This study contributes to understanding the biological treatment of nitroaromatics, offering insights into environmental pollution mitigation strategies and the development of efficient bioremediation techniques.

## INTRODUCTION

Toxic compounds can be categorized into natural compounds, by-products of metabolism, xenobiotic compounds, products of human activities, etc. Due to their complex structures, xenobiotics often exhibit low biodegradability, posing significant environmental challenges. Since these compounds have a very recent origin, organisms that can degrade these compounds are limited (Rieger *et al*., 2002). 2,4-DNP is a di-nitro aromatic organic compound with the chemical formula HOC_6_H_3_(NO_2_)_2_. It is a refractory pollutant released into the environment from industrial effluents, spill-outs, or through the degradation of pesticides containing 2,4-DNP. According to the US Environment Protection Agency, it is a priority pollutant whose concentration in natural water sources should be limited to 10mg/L (Long *et al*.,2003).

Several physical and chemical methods are known for treating nitrophenol pollutants, but they are not safe and cost-effective. Energy-intensive chemical treatments such as incineration may be too expensive at low concentrations or may cause other environmental problems such as NO_x_ emissions. Other relevant biological procedures which are safe and cost-effective are being investigated for the purpose. Some microbes have developed mechanisms to degrade toxic compounds through reductive pathways despite the structural rigidity conferred by the delocalization of electrons in aromatic rings (Esteve-Núñez *et al.,* 2001; Ramos *et al*., 2005). However, safe and inexpensive strategies to reduce nitro phenolics is of concern and an important area of study.

Cellular proteolysis is used to remove aberrant and misfolded proteins and restrict the quantity and duration of essential regulatory proteins. In prokaryotes, energy-dependent protein degradation is controlled majorly by two classes of proteases: Cytoplasmic and membrane-bound. The major cytoplasmic proteases are Lon (serine protease), Clp(serine protease) and HS1UV(threonine protease) whereas FtsH(metalloprotease),EcfE(zinc metalloprotease) and DegS(serine protease) are membrane-bound (Chandu & Nandi, 2004; Neuwald *et. al*., 1999; Lngklotz *et al*., 2012; Sohn *et al*., 2007). The Lon protease is a member of the AAA^+^ family of proteins (<u>A</u>TPases <u>A</u>ssociated with various cellular <u>A</u>ctivities) with three domains: amino-terminal, central domain and carboxy-terminal domain. The amino-terminal recognizes and binds to substrates, the central ATPase domain contains an ATP binding motif whereas the carboxy-terminus contains the catalytic serine-lysine dyad for proteolysis (H. Fischer & R. Glockshuber, 1994; W. Ebel *et.al*,1999; Nomura *et.al*, 2004).

Worldwide research is being conducted on the use of bioremediation to effectively decontaminate nitroaromatics. Particularly of interest to biotechnologists are enzymes that facilitate the reduction process. The possible role of Lon protease in the metabolism of 2,4- DNP is an unexplored area of research. In this study, we aim to decipher the roles of *E. coli*- encoded Lon protease in the metabolism of 2,4-DNP. The study highlights Lon protease’s pivotal role in converting xenobiotic nitroaromatic 2,4-DNP by *E. coli.* The enhanced conversion observed in the Δ*lon* strain highlights the influence of Lon on nitro reductase activity, likely mediated through the MarA regulatory system. This work provides valuable insights into the biological pathways involved in nitroaromatic compound degradation by characterising the distinct conversion products and elucidating the regulatory mechanisms.

## Results

### UV-visible spectral pattern of conversion products of DNP differs in WT and Δ*lon* strains 18 hr post-treatment

During our studies with uncouplers (Verma *et al*., 2024), we observed the formation of a reddish-brown colour in WT and Δ*lon* strain with 2,4-DNP and the intensity of colour in the Δ*lon* strain was higher. WT and Δ*lon* strain incubated with 2,4-DNP (yellow colour) started showing change in the colour of supernatant from 12 hr post-treatment, where both the supernatants showed dark orange colour. However, from the 18^th^ hour post-treatment, the supernatant of Δ*lon* strain showed a change to a reddish-brown colour, and the colour was visually more intense in the Δ*lon* strain. These observations were investigated in greater detail and bacterial cultures were grown in 5ml Minimal Media (MM) with and without 0.5mM 2,4- DNP for 6, 12, 18 and 24 hr and cells were pelleted down at 10,000xg for 10 mins. The supernatant was collected and filtered, and UV-vis spectra of the supernatant were recorded in a UV-vis spectrophotometer. The reduction in the concentration of 2,4-DNP during the conversion was quantified using the formula (1), and the ratios are plotted in a graph (Fig: 1, SI Fig: 1). The spectral pattern of 2,4-DNP was similar to the previously reported pattern (Srinivasan *et al*., 2012; Khabarov *et al*.,2012), with a peak maximum of 364 nm. There was a change in the peak maxima for the new compounds formed when 2,4-DNP was inoculated with the Δ*lon* strains from 12 hr post-treatment (SI fig 1); however, a significant difference in the peak pattern of the supernatant obtained from the two strains under investigation compared to 2,4-DNP spectra was observed at 18 hr post-treatment (Fig 1A). A reduction in absorption of 2,4-DNP peak maximum could be observed in the WT supernatant, and a red shift could be observed in the supernatant from the Δ*lon* strain. The WT supernatant showed a peak of 364 nm, whereas the mutant supernatant gave a peak of 394 nm.

**Figure 1.**
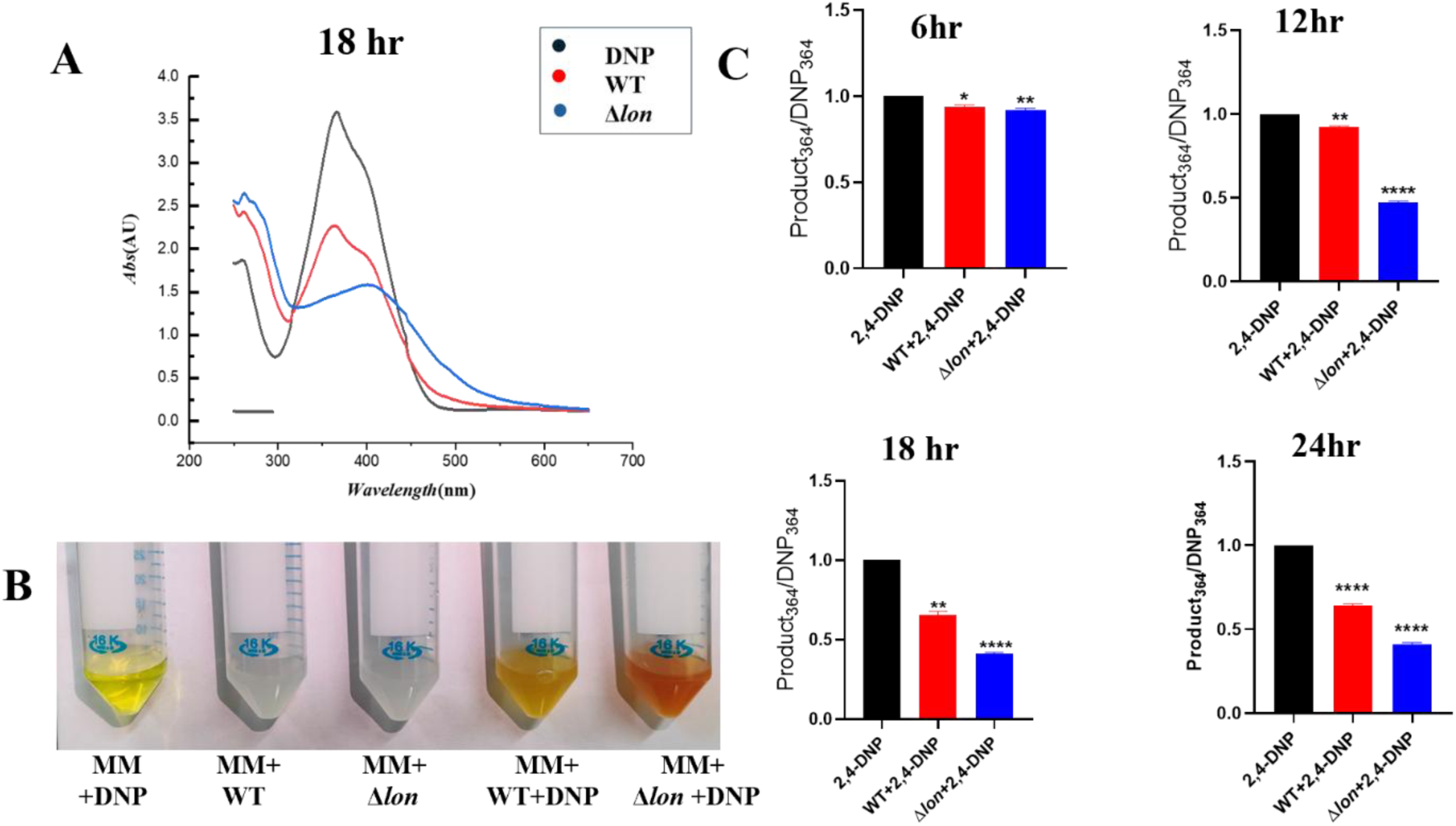
A reddish-brown-coloured compound is produced in higher amounts in the Δ*lon* strain grown in minimal media in the presence of 2,4-DNP. *E. coli* MG1655 WT and Δ*lon* strains were cultured for 18 hr at 37°C and 160 rpm in the presence of 0.5mM of 2, 4-DNP. Culture supernatants were collected from bacterial culture grown in the absence and presence of 0.5 mM 2,4-DNP. (A) UV-visible spectrum of the supernatant after diluting 1:2 (B) Representative image showing conversion product formed post 18hr of treatment. (C) Quantification of the spectrum was performed for 6^th^,12^th^,18^th^ and 24^th^ hr. The data are representative of three independent experiments plotted as mean ± SD. Statistical analysis was performed using two-way ANOVA and one-way ANOVA for (C), where * indicates p<0.05.

### The higher conversion product seen in the Δ*lon* strain is not due to the higher growth of the mutant in the presence of 2,4-DNP

Overall, it appeared that the Δ*lon* strain could convert 2,4-DNP (yellow) to a reddish-brown colour in the growth media faster than the WT strain. The effect could be due to the higher growth of Δ*lon* strain in the presence of 2,4-DNP. Therefore, we decided to study the growth kinetics of both strains in the presence and absence of 2,4-DNP in MM. The bacterial strains were grown in 50ml MM with and without 0.5mM 2,4-DNP, and the optical density at 600 nm (O.D._600nm_) of bacterial culture was measured at regular intervals from 0 to 24 hr. Colony-forming unit/ml (CFU/ml) was calculated by spread plating 50 μl of different dilutions of the bacterial culture collected at regular intervals in LB agar plates, followed by overnight growth and counting the colonies using a colony counter. A dose screening was done to find the optimal dose for the growth experiment with 2,4-DNP in MM, and 0.5mM was found to be the optimal dose to study the growth kinetics (SI Fig 2A). The growth rate of the Δ*lon* strain was less than WT in the presence of 2,4-DNP in MM from 3 hr post-treatment (SI Fig 2B,C). It could be inferred that more conversion product seen in Δ*lon* was not due to more growth of the knock- out strain. Thus, it is likely that the difference in conversion observed was due to changes in the metabolism of 2,4-DNP in the absence of Lon protease.

**Figure 2.**
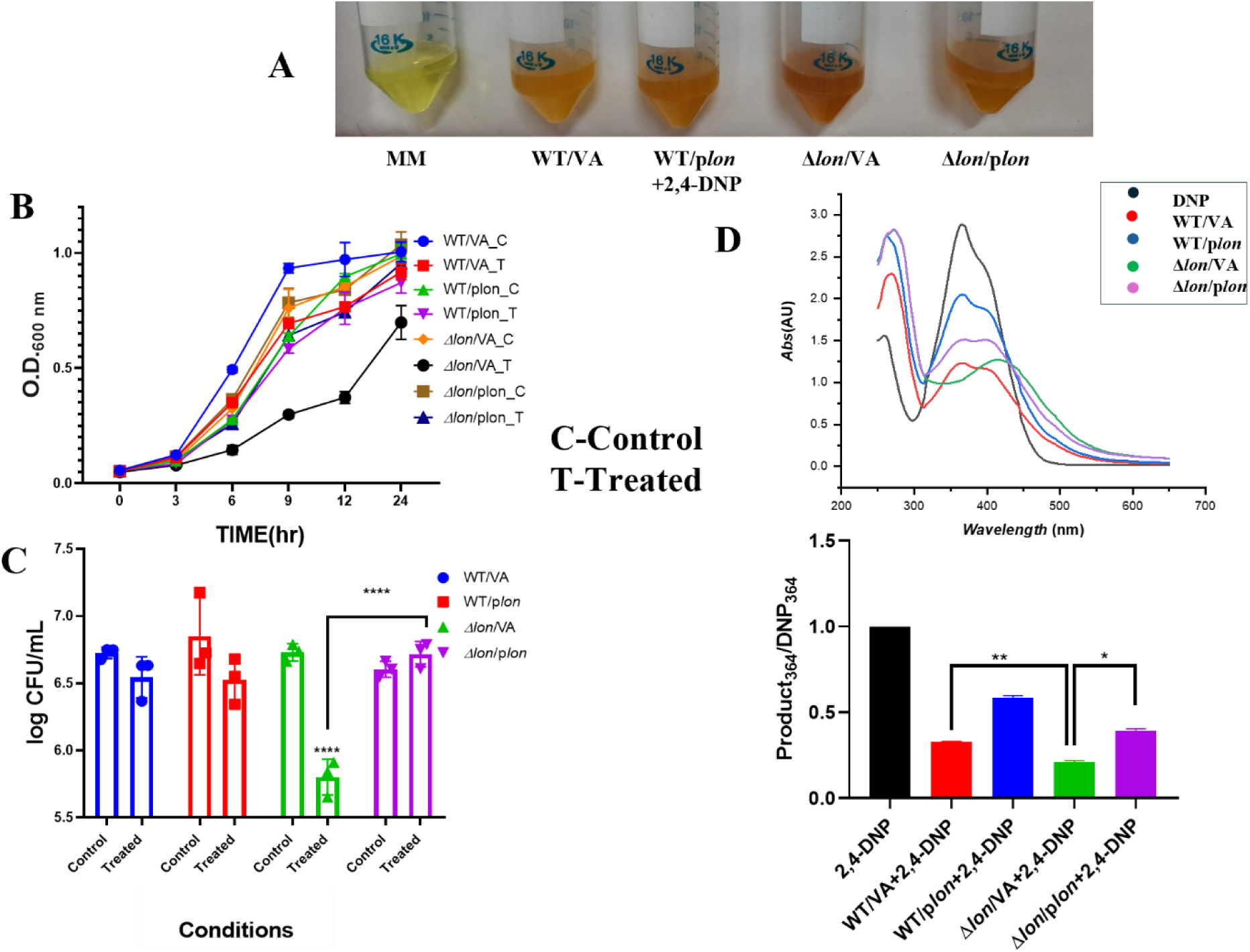
Trans *c*omplementation of *lon* increases the growth of Δ*lon* in the presence of 2,4-DNP but lowers the amount of reddish-brown coloured compound. *E. coli* WT /VA, WT /p*lon,* Δ*lon/*VA, and Δ*lon/*p*lon* were cultured in the presence of 0.5mM 2, 4-DNP for a period of 18 hr at 37°C and 160 rpm. (A)Representative image showing conversion product formed post 18hr of treatment; (B) Growth of the strains during exposure to 0.5 mM 2,4-DNP is shown; (C) CFU count is shown 6 hr post-exposure; (D) UV-visible spectrum of the supernatant after diluting 1:2 with a graph showing quantification of the spectrum. C represents the control group, and T represents the treated group. The data are representative of three independent experiments plotted as mean ± SD. Statistical analysis was performed using two-way ANOVA and one-way ANOVA for (D), where * indicates p<0.05.

### Trans complementation of *lon* increases the growth of Δ*lon* but reduces the amount of reddish-brown colour in the presence of 2,4-DNP

Trans-complementation was carried out to confirm the *lon* dependency of the phenotypes observed with the strain. Growth and spectral patterns of the growth supernatant of strains transformed with the empty vector and plasmid with the gene of interest were investigated. Trans complementation of *lon* partially rescued the peak shift observed in Δ*lon*/VA (plasmid vector transformed strain) to the WT spectral pattern discussed in the previous result (Fig 2).

Quantifying the spectra further confirms the partial rescue of DNP drop-in Δ*lon*/p*lon* (plasmid containing *lon* transformed strain). The growth reduction induced by 2,4-DNP in Δ*lon*/VA was also rescued in Δ*lon*/p*lon* (Fig 2B&C). WT/p*lon,* the over-expression system of *lon* showed a lesser drop in 2,4-DNP (Fig 2D). All these confirm that the growth phenotype and the spectral pattern observed is *lon*-dependent.

### The conversion of 2,4-DNP into the reddish-brown colour is MarA-dependent

The Lon protease has various substrates and several strains lacking many of these substrates are already available in our laboratory (Verma *et al*., 2024). Single knock-outs of *marA, rob* and *soxS* were screened for their ability to convert 2,4-DNP to the reddish-brown colour (SI Fig 3). The Δ*rob* and Δ*soxS* strains converted 2,4-DNP to the reddish-brown colour although some changes were seen in the Δ*rob* strain compared to WT (SI Fig 3C). However, the Δ*marA* strain completely failed to convert 2,4-DNP to the reddish colour. Also, the Δ*lon*Δ*marA* strain failed to convert 2,4-DNP to the reddish-brown colour which showed the complete dependence of the conversion phenomenon on *marA* (Fig 3). To further confirm the phenomenon, trans complementation of *marA* was generated and the WT/VA and Δ*marA*/VA strains showed the phenotypes of WT and Δ*marA*, respectively, in terms of conversion and growth. WT/p*marA* showed higher conversion, depicting the role of higher amounts of *marA* in a higher substrate compound conversion. Δ*marA*/p*marA* rescued the conversion and growth reduction phenotype observed in Δ*marA* compared to WT phenotype (Fig 4). All these results confirm that the conversion of 2,4-DNP to the reddish-brown colour is entirely *marA-*dependent.

**Figure 3.**
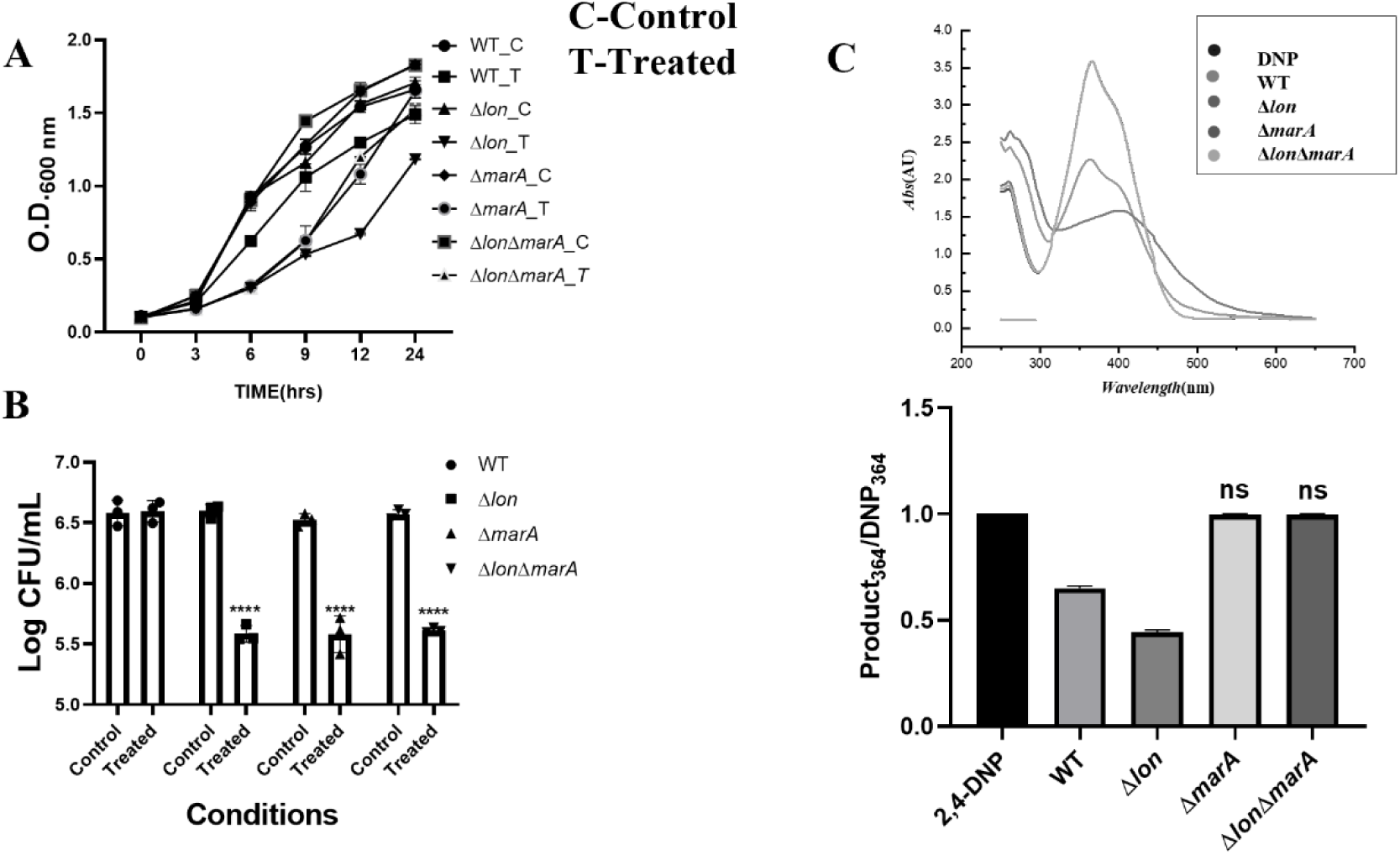
The *marA* mutant fails to convert 2,4-DNP to the reddish-brown coloured compound. *E. coli* WT, Δ*lon*, Δ*marA* and Δ*lon* Δ*marA* were cultured in the presence of 0.5mM 2, 4-DNP for a period of 18 hr at 37°C and 160 rpm. (A) The growth of the strains during exposure to 0.5 mM 2,4-DNP is shown; (B) CFU count is 6 hr post-exposure; (C) UV-visible spectrum of the supernatant after diluting 1:2 with a graph showing quantification of the spectrum. C represents the control group, and T represents the treated group. The data are representative of three independent experiments plotted as mean ± SD. Statistical analysis was performed using two-way ANOVA for (C) and one-way ANOVA for (D), where * indicates p<0.05.

**Figure 4.**
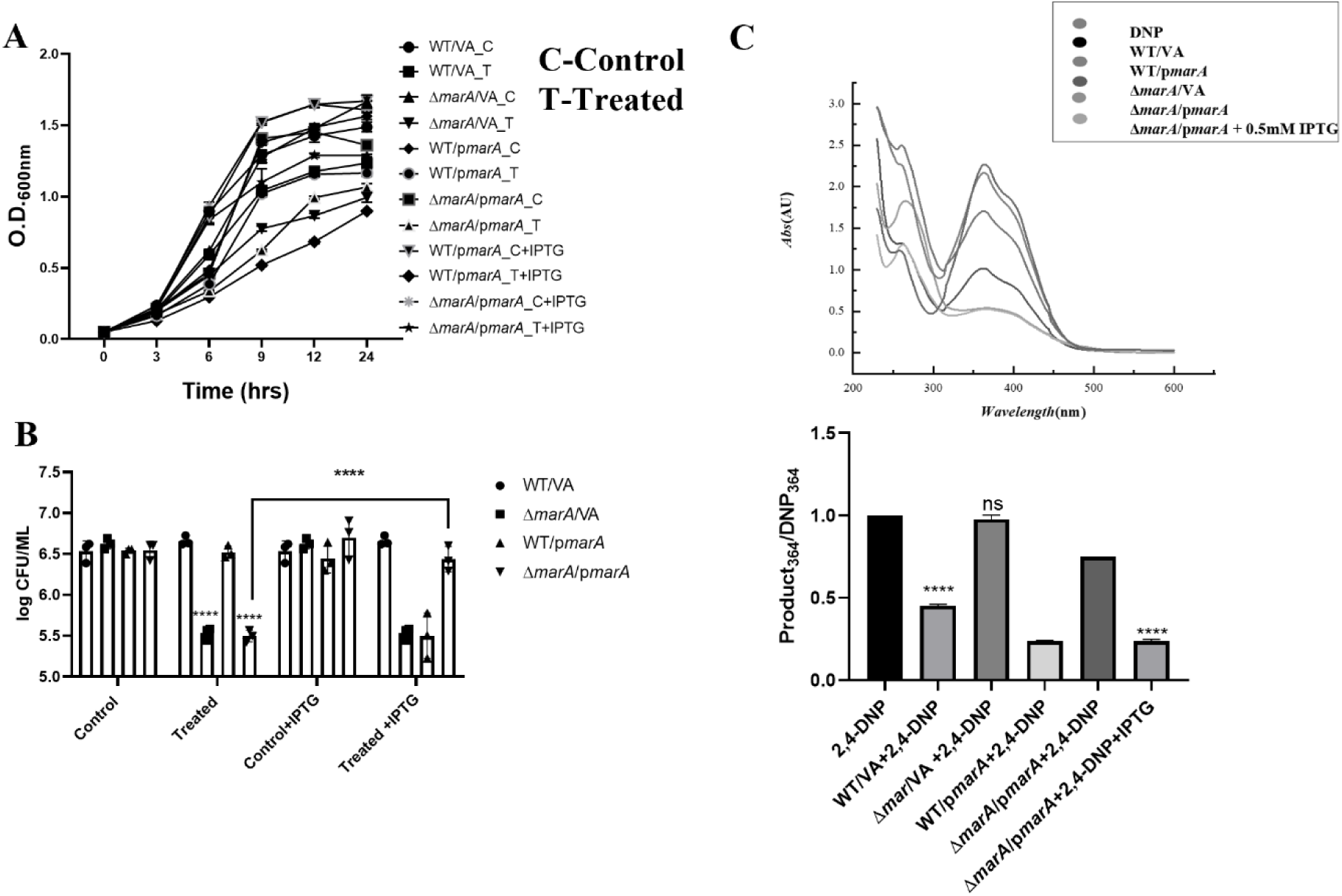
Trans-complementation confirms the *marA* dependency of 2,4-DNP metabolism. *E. coli* WT/VA, WT/ /p*marA,* Δ*marA*/VA, and Δ*marA*/p*marA* were cultured in the presence of 0.5mM 2, 4-DNP for a period of 18 hr at 37°C and 160 rpm. (A) The growth of the strains during exposure to 0.5 mM 2,4-DNP is shown; (B) CFU count is shown 6 hr post-exposure;(C) The UV-visible spectrum of the supernatant after diluting 1:2 with a graph showing quantification of the spectrum. C represents the control group, and T represents the treated group The data are representative of three independent experiments plotted as mean ± SD. Statistical analysis was performed using two-way ANOVA C) and one-way ANOVA for (D), where * indicates p<0.05.

### The nitroreductase deletion mutants failed to convert 2,4-DNP to the reddish-brown colour

Both *nfsA* and *nfsB* (nitroreductase genes) have a MarA box (SI Fig 4), which induces the expression of the two genes when the *mar* operon is derepressed (Barbosa TM & Levy SB, 2000). Since the roles of *lon* and *marA* were confirmed, we evaluated the expression of genes encoding these nitroreductases and their association with *marA* in the context of 2,4-DNP metabolism. The levels of *marA*, *nfsA* and *nfsB* were upregulated with 2,4-DNP treatment in both WT and Δ*lon* strains; however, the upregulation was not observed in Δ*marA* post-treatment with 2,4-DNP (Fig 5). These results establish the relation of *marA* with the induction of nitroreductase genes in the metabolism of 2,4-DNP. To further understand the possible functional roles of the nitroreductase genes in the metabolism, *nfsA* and *nfsB* knock-outs were screened. Δ*nfsA* and Δ*nfsB* showed slight reduction in the amounts of 2,4-DNP in the supernatant compared to DNP in MM control (Fig 5 D, E), demonstrating that both genes are involved in the conversion of 2,4-DNP. These results confirmed the roles of *marA* dependence on the metabolism and gain insights into the importance of both nitroreductases for conversion.

**Fig 5.**
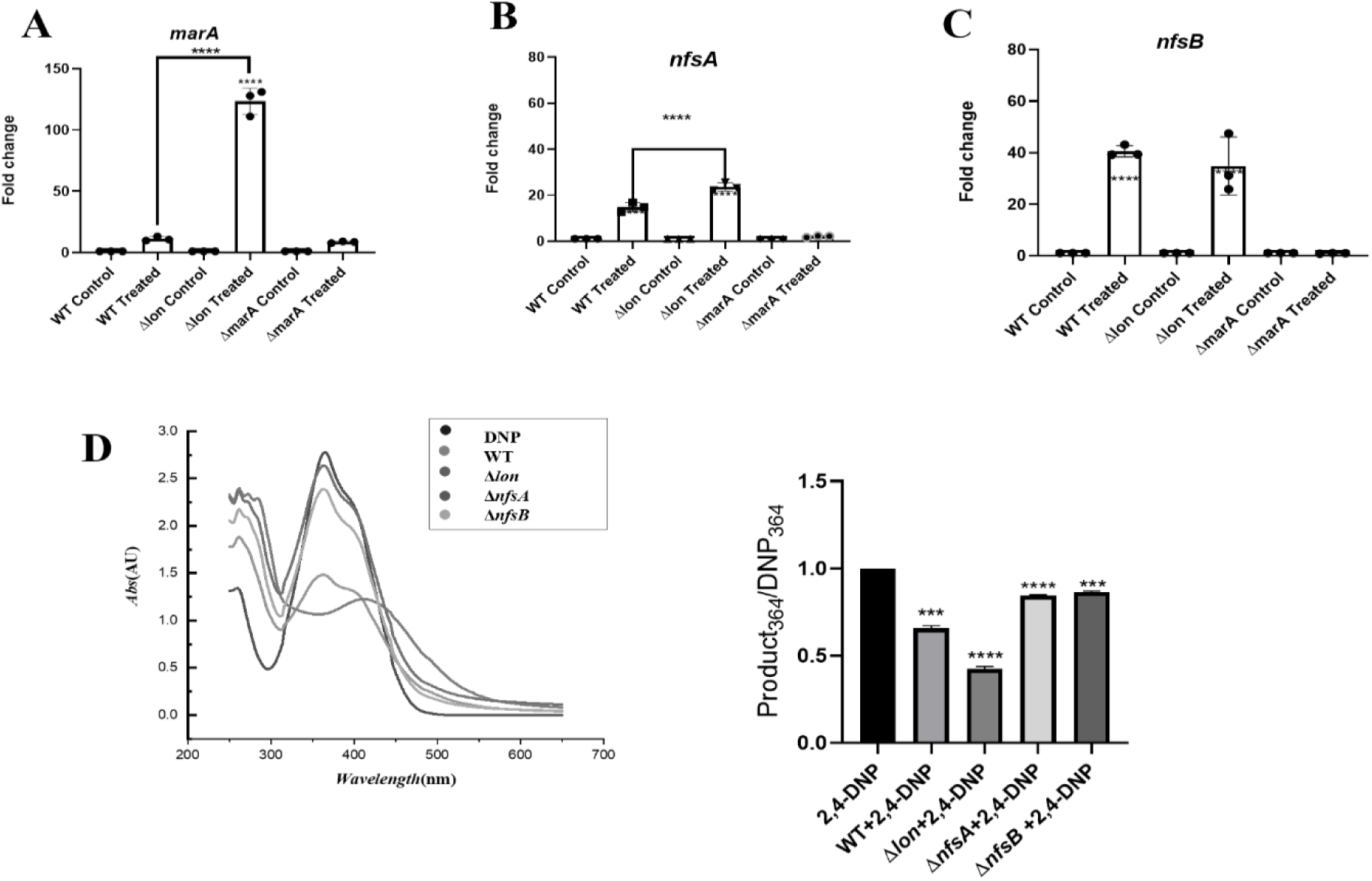
2,4-DNP induces the expression of *marA*, *nfsA* and *nfsB* in both WT and Δ*lon* strains but not in the Δ*marA strain*. The cells were grown for 3 hours at 37^○^C, and then 2, 4-DNP was added for the treated condition; cells were harvested 3 hr post-treatment and processed for RNA isolation. Fold changes in transcript levels of (A) *marA*, (B) *nfsA*, and (C) *nfsB* in WT, Δ*lon* and Δ*marA* strains were determined by quantitative real-time PCR. *gapA* was used as the reference gene. *E. coli* WT, Δ*lon*, Δ*nfsA* and Δ*nfsB* were cultured in the presence of 0.5mM 2, 4-DNP for a period of 18 hr at 37°C and 160 rpm. (D) UV-visible spectrum of the supernatant after diluting 1:2 with and graph showing quantification of spectrum. The data is representative of three independent experiments with mean ± SD where * indicates P<0.05. Statistical analysis was performed for each strain relative to its untreated control. Comparison between the strains is indicated wherever significant.

### 4-Amino-2-nitrophenol is the compound corresponding to the peak m/z=153.03 in the LC-MS spectrum

To identify the conversion product, we analysed the supernatant using LC-MS and chose this strategy due to its high accuracy. The chromatogram obtained from LC-MS samples was overlapped, and the common and unique peaks were identified and marked. The intensity of the peaks was also analysed to understand the conversion product of 2,4-DNP and other metabolites being released into the growth suspension. The suspension of 2,4-DNP clearly showed a single peak corresponding to an m/z value of 183.0045, which, upon analysis and fragment search, was confirmed to be from 2,4-DNP (Fig 6 A, E). This served as a control and validated the experimental flow and the peak intensity was 2x10^7^. The chromatogram obtained from the growth suspension of the WT strain showed 3 major peaks at m/z = 183.0045, 153.0310 and 329. 0550 (Fig 6A). The first peak was identified as corresponding to 2,4-DNP but with a lower intensity of 1.2x10^7,^ as 2,4-DNP was being metabolized by the bacteria during growth. The second peak of m/z=153.0310 had two hits with a final score of 1.0, which were 4-Amino-2-nitrophenol and 2-Amino 4-nitrophenol and the peak intensity was 0.3x10^7^. The Δ*lon* strain showed a few major peaks, out of which two peaks corresponding to m/z= 183.0045 and 153.031 could be identified as 2,4-DNP and 2,4-ANP, respectively (Fig 6C). The peak intensities of 2,4-DNP and 2,4-ANP were around 0.1x10^7^ and 0.7x10^7^ respectively. Since LC-MS relies solely on a molecule’s mass/charge ratio, isomeric forms cannot be differentiated. Since the nitro group in the meta position of 2,4-DNP cannot undergo reduction due to the electron-withdrawing nature of the hydroxyl group in the ortho position, the compound was confirmed to be 4-amino-2-nitrophenol (Fig 6D). The third peak of m/z=329.0550 did not give any hits from the KEGG database. Since the identified compound is reddish-brown, the conversion product can be confirmed as 4-amino-2-nitrophenol.

**Figure 6.**
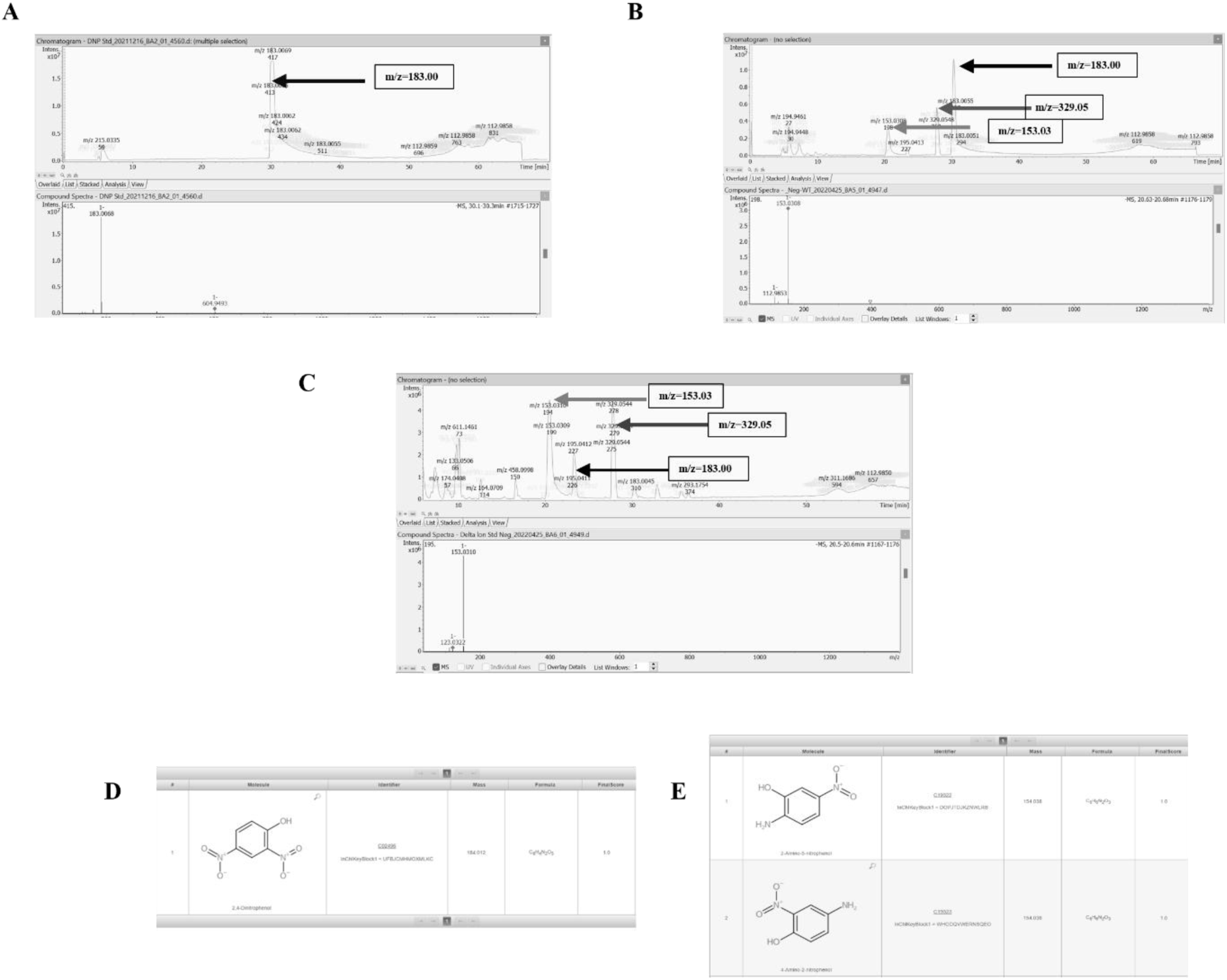
A compound corresponding to the peak m/z=153.03 is confirmed to be 4-amino-2-nitrophenol. The cells were grown for 18 hours at 37^○^C in the presence of 2, 4-DNP, supernatant was collected and processed for LC-MS. (A) The suspension of 2,4-DNP clearly showed a single peak designating to an m/z value of 183.0045, which upon (D) analysis and fragment search was confirmed to be from 2,4-DNP; (B) WT strain showed 3 major peaks at m/z = 183.0045, 153.0310 and 329. 0550 and (E) peak of m/z=153.0310 had two hits with a final score of 1.0 they were 4-Amino-2-nitrophenol and 2-Amino-4-nitrophenol; (C) Δ*lon* strain showed many major peaks out of which two peaks corresponding to m/z= 183.0045 and 153.031 could be identified as 2,4-DNP and 2,4-ANP respectively.

### 4,2-ANP induces phenotypic antibiotic resistance

A previous study from our lab has shown the ability of 2,4-DNP to induce antibiotic resistance in a *marA*-dependent manner (Verma *et al*., 2024). In the current study, we found that 2,4-DNP is reduced to 4,2-ANP by *E. coli*. In a continuous culture with a regular flow of nutrients, the conversion product can affect the metabolism and be a possible inducer of phenotypic antibiotic resistance. To answer the above questions, we screened WT with different doses of 4,2-ANP and compared the growth with 2,4-DNP in LB. As previously reported, 2,4-DNP reduced the growth of the WT strains (Verma *et al*., 2024); however, no significant lowering of growth was observed by 4,2-ANP (Fig 7A). 2,4-ANP is a commonly used non-carcinogenic compound in hair colour dyes and is considered safe for humans (Burnett *et al*., 2009), although the effect on AMR remains unexplored. Broth dilution MIC and growth assay was used to test the ability of 4,2-ANP to induce antibiotic resistance against ciprofloxacin and tetracycline and compared it with DNP. Both compounds induced antibiotic resistance against ciprofloxacin and tetracycline.2,4-DNP was more potent in inducing resistance against ciprofloxacin whereas 4,2-ANP was more potent in case of tetracycline resistance induction (Fig 7 C, D).

**Fig 7.**
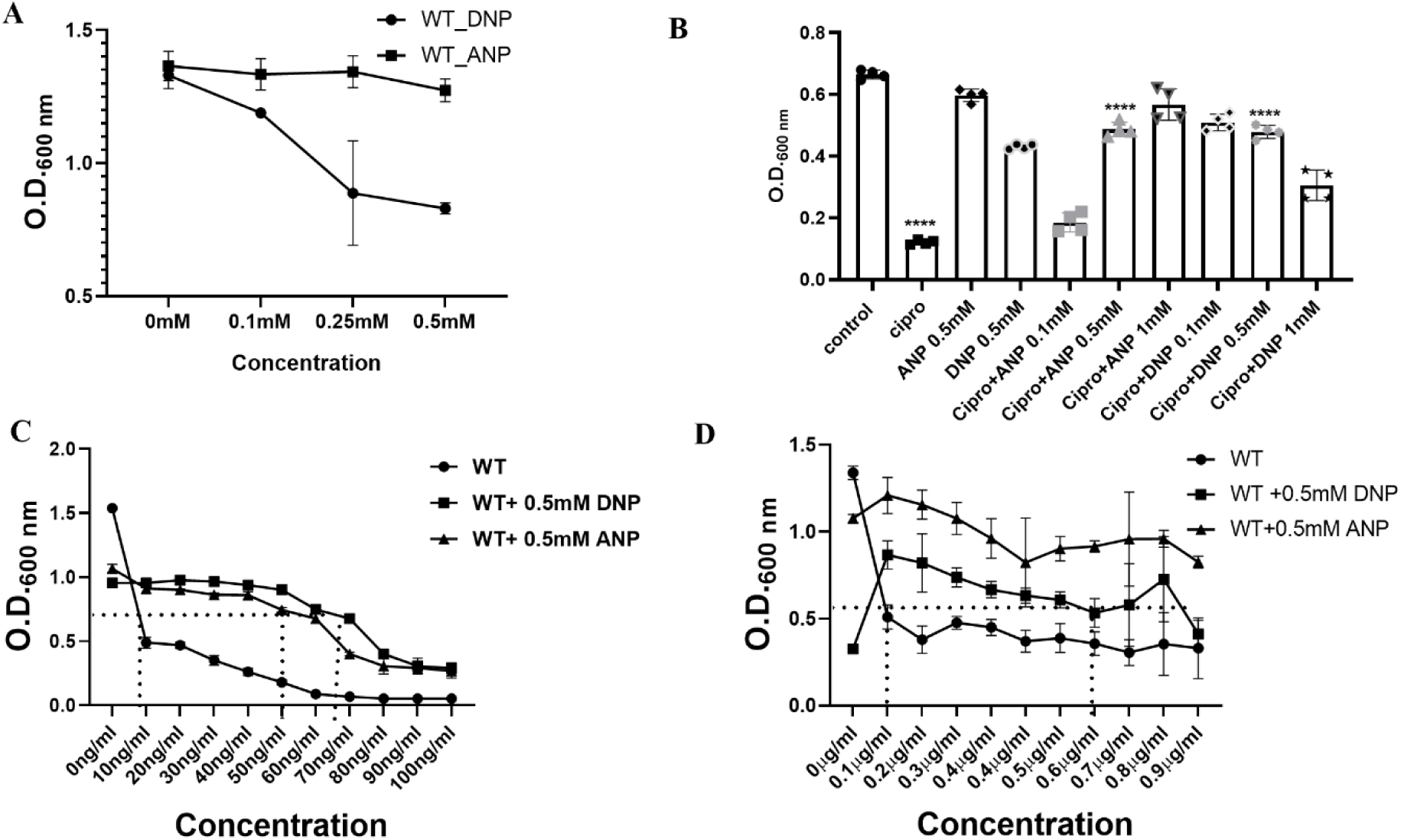
4,2-ANP induces phenotypic antibiotic resistance against ciprofloxacin. *E. coli* WT strain was cultured with concentrations of 2, 4-ANP and 4-DNP for 6 hr at 37°C and 160 rpm in LB.; (A) The growth of WT strain under different concentrations of 2,4-DNP and 4,2-ANP. (B) The growth of strains under 20ng/ml of ciprofloxacin and in combination with different concentrations of 2,4-DNP and 4,2-ANP;MIC broth dilution assay with (C) ciprofloxacin and (D) Tetracycline was performed, and data is represented as line plot. The data are representative of at least three independent experiments plotted as mean ± SD. * indicates P<0.05. Statistical analysis was performed for each strain relative to its untreated control. Comparison between the strains is indicated wherever significant.

**Fig 8.**
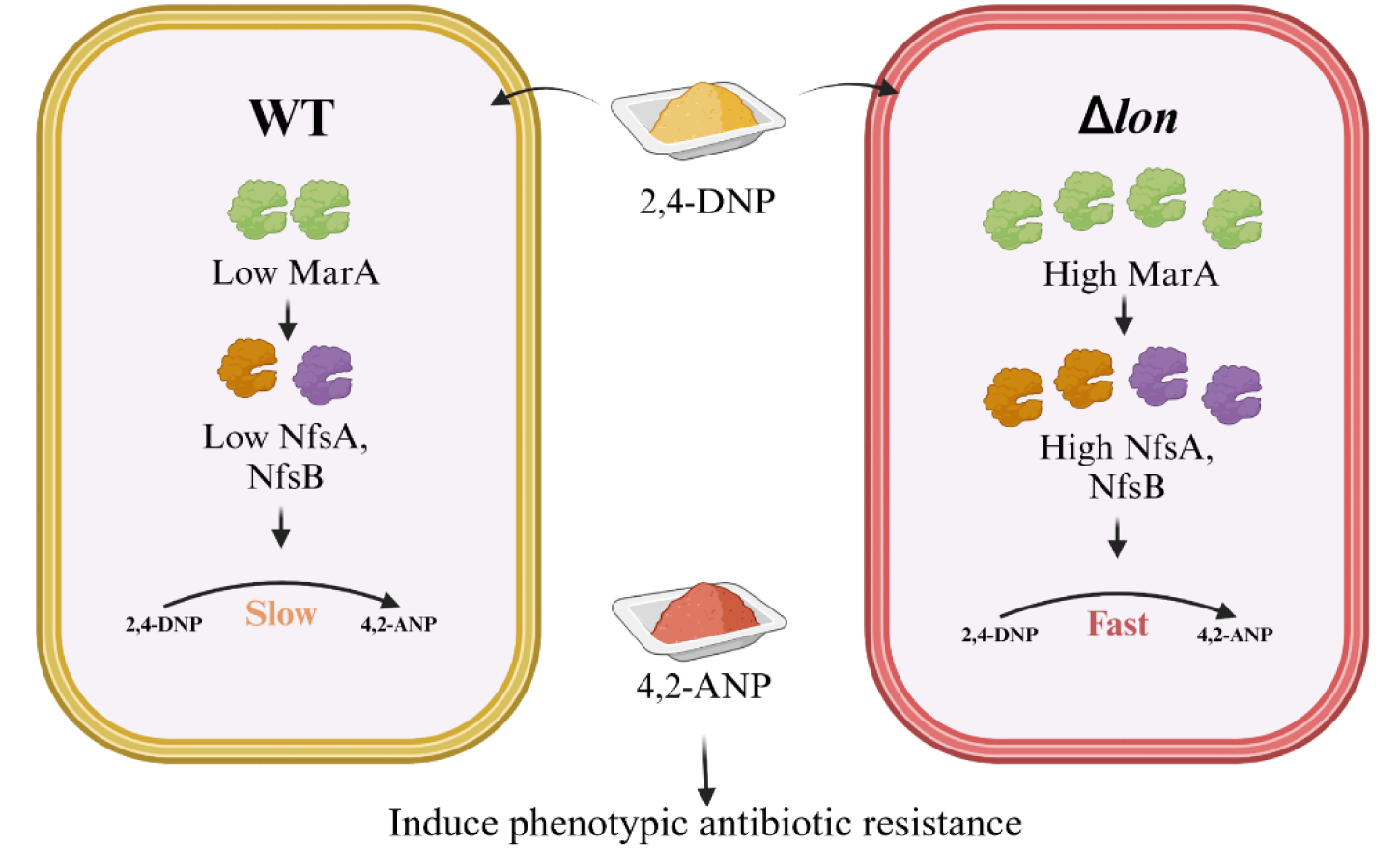
A model representing the roles of Lon protease and its substrate MarA in the generation of 4,2-ANP, which induces phenotypic antibiotic resistance. The role of Lon protease in degrading the excess amount of MarA, which induces nitroreductases NfsA and NfsB in the metabolism of 2,4-DNP, is shown. The study identifies 4,2-ANP as a novel inducer of phenotypic antibiotic resistance

## Discussions and Conclusions

This work details the roles of Lon protease and the MarA transcription in the conversion of 2,4-DNP to 4,2-ANP and the identification of the latter as an inducer of phenotypic antibiotic resistance. Previously published reports from our group have shown that the *lon* deletion strain is sensitive to DNP stress due to higher amounts of *marA* in nutrient-rich media (Verma *et al*., 2024). A similar phenotype was noticed when Δ*lon* was subjected to different doses of 2,4- DNP in minimal media. A more intense reddish-brown colour was observed in the tubes from the 18^th^ hour, even with lesser growth than WT (Fig. 1). UV-visible spectroscopy is a non-invasive analytical technique that measures the amount of UV or visible light absorbed or transmitted by a matter. UV-Vis spectroscopy was used to analyse the change in spectral pattern and identification of the conversion product (Nilapwar,2011). A spectrum with a peak maximum at 364nm was observed for 2,4-DNP, similar to previous reports (Srinivasan *et al*., 2012). The WT supernatant showed similar peak maxima with lesser absorbance, which indicated the metabolism of the substrate, i.e., 2,4-DNP. However, the Δ*lon* supernatant showed a new spectral pattern with a bathochromic shift and a peak maximum at around 410 nm (Fig 1). Although a change in peak pattern was observed, the final product could not be concluded with the spectral patterns and peak maxima due to the unavailability of a database. It is also possible that the peak maxima and pattern couldn’t match the available data due to a mixture of compounds in the media hindering the recording of a clean compound spectrum. The growth reduction phenotype and the conversion pattern in Δ*lon* were also trans- complementable (Fig 2).

The bathochromic shift indicates a reduction reaction, which pointed out the involvement of nitroreductases and intrigued us to study the possible interaction of Lon and nitroreductases in the context of 2,4-DNP metabolism. Nitro reductases in bacteria are flavoenzymes with NADH as an electron donor that catalyses the reduction of the nitro group in flavins, quinones, nitro heterocyclic, and nitroaromatic compounds. (de Oliveira *et al*., 2007). These enzymes have been widely studied due to their potential application in bioremediation and biomedicine (Spain, 1995). Although most nitro reductases have been purified and their biochemical properties characterized, their physiological roles remain obscure. Two types of bacterial nitro reductases have been described based on their response to oxygen:1) oxygen-insensitive or type I nitro and 2) oxygen-sensitive or type II nitro reductases (Bryant *et al*., 1981; Mason & Holtzman, 1975). Their phylogenetic analysis suggests that the type I nitroreductase can be classified into two families, NfsA and NfsB, dependent on NADPH and NADPH or NADH, respectively, as the electron donor. In *E. coli,* the *nfsA* and *nfsB* genes code for 27 and 24 kDa proteins termed major and minor nitro reductases, dissimilar in their amino acid sequence level (Bryant *et al*., 1981; Zenno *et al*., 1996; Whiteway *et al*., 1998). Bacterial nitro reductases are believed to be regulated by *marRAB* and *soxRS* operon systems (Barbosa TM & Levy SB, 2000; Barbosa TM & Levy SB, 2002). Numerous studies have shown that MarA, Rob and SoxS are closely related and can bind to a degenerate sequence known as the *mar* box, overlapping the genes they regulate (Duval *et al*.,2013). The nitroreductase genes also have a *mar* box in their upstream sequence (Barbosa and Levy, 2002). Similar to NaSal, 2,4-DNP can also bind to MarR and initiate the transcription of *marA* (Heuveling *et al*.,2008). It is also known that Lon protease can maintain the homeostasis of these three transcription factors, and the absence of Lon can lead to higher amounts of transcription factors in the cell (Bhaskarala *et al*.,2016). Therefore, we screened these genes (*marA*, *rob* and *soxS*) for their ability to convert DNP and found that the conversion was entirely *marA-*dependent since Δ*marA* cells failed to show any reduction in the intensity of 2,4-DNP (Fig 3). Also, the Δ*lon*Δ*marA* strain showed no conversion and spectral pattern overlapped with that of 2,4-DNP alone (Fig 3), like Δ*marA,* confirming that MarA was responsible for the phenotype observed in the strain lacking Lon, i.e., higher levels of MarA in Δ*lon* leads to the higher conversion of 2,4-DNP. Additionally, qPCR showed the upregulation of nitroreductase genes (*nfsA*, *nfsB*) in both WT and Δ*lon* upon treatment with 2,4-DNP, and the upregulation was higher in Δ*lon*. However, no upregulation of nitroreductase genes was observed in Δ*marA* cells upon treatment with 2,4-DNP (Fig 5 A-C). These results confirm the interplay between Lon, MarA, *nfsA* and *nfsB* upon treatment with 2,4-DNP. Since both the nitroreductases showed upregulation, we could not conclude the key nitroreductase involved in the metabolism of 2,4-DNP. To resolve this, we utilised the knock-out strains of *nfsA* and *nfsB* and screened them for their ability to convert 2,4-DNP. To our surprise, both knockouts failed to show conversion of 2,4-DNP (Fig 5 D).

A high throughput assay was used to confirm the compound formed in the growth suspension (Yu S *et al*., 2024). LC-MS was carried out from the growth suspension, and the presence of 4,2-ANP was confirmed in both the WT and Δ*lon* strains. There were many other compound peaks in the LC-MS spectrum, but most remained unresolved, mainly carbohydrate or fatty acid molecules and mostly insignificant for our study. The concentration of 4,2-ANP was found to be higher in the Δ*lon* strain together with reduction in the concentrations of 2,4- DNP (Fig 6). 4,2-Amino nitrophenol has been previously studied in various aspects, including safety for usage in hair colour dyes, induction of mutation in *Salmonella* Typhimurium, percutaneous absorption in human and monkey skin and effects on the metabolism of rats (Burnett *et al*.,2002; Shahin *et al*.,1982; Bronaugh RL and Maibach HI,1985; Cameron M A, 1958). However, the effect on the metabolism of *E.coli* and induction of phenotypic antibiotic resistance by 4,2-ANP remains an unexplored area. Compared with the substrate molecule 2,4-DNP, 4,2-ANP was less potent in reducing growth and induction of resistance against ciprofloxacin, but it was more potent than 2,4-DNP in inducing resistance against tetracycline (Fig 7). This is a novel finding, and the mechanism by which resistance is induced needs to be explored further.

Nitroaromatic compounds like 4,2-ANP are widely used in industrial applications, including dye production. The findings suggest that their environmental presence may inadvertently influence microbial communities, impacting ecosystems and potentially human health. This study adds to the limited body of knowledge regarding the metabolic and phenotypic effects of nitroaromatic compounds on bacteria. It introduces a new perspective on how such compounds interact with microbial systems, paving the way for further research into metabolic pathways and regulatory mechanisms. These findings have broader implications for bioremediation, suggesting that strains lacking Lon protease or engineered to enhance nitroreductase activity could be optimised for environmental applications. These results contribute to our understanding of the microbial metabolism of toxic compounds and open avenues for sustainable solutions to industrial pollution through targeted biotechnological innovations. Also, this study emphasises the need to carefully assess chemical pollutants and their unintended consequences on microbial resistance. Ultimately, this study lays the foundation for developing efficient microbial systems for detoxifying and bioremediation harmful environmental pollutants.

## Materials &Methods

### Bacterial strains and growth conditions

The bacterial strains used in this study are listed in Table 1. All the cultures were grown in Luria Broth (LB) comprising Tryptone (10 g/L; HiMedia, Mumbai, India), NaCl (10 g/L; HiMedia) and yeast extract (5 g/L; HiMedia) at 37°C and 160 rpm. Overnight grown cultures obtained from a single colony of the strains served as per-inoculum for all experiments. Antibiotics were used at 100 μg/ml ampicillin, 30 μg/ml chloramphenicol, and 50 μg/ml kanamycin (HiMedia). Cultures for experiments were carried out in Minimal media (MM) containing 64 g/l Na_2_HPO_4_•7H_2_O, 15 g/l KH_2_PO_4_, 2.5 g/l NaCl, 5.0 g/l NH_4_Cl. The optical density (O.D.) of all the strains was normalised to O.D. 2, which corresponds to 10^9^ CFU/ml, before all experiments, and 0.2% and 1% of these O.D. 2 cultures were used as the starting inoculum for all experiments in LB and MM respectively for all experiments. A UV-visible spectrophotometer (Tecan, Männedorf, Switzerland) determined growth by measuring the O.D._600 nm_.

**Table 1:** List of strains used in the study.

| Sl.No | Strain | Genotype/feature | Reference |
| --- | --- | --- | --- |
| 1 | WT MG1655 | $\lambda$, rph-1 | Guyer *et al.* (1981) |
| 2 | $\Delta lon$ | MG1655, *lon* : : *cat* | Bhaskarla *et al.* (2016) |
| 3 | $\Delta marA$ | MG1655, *marA* : : *kan* | Baba *et al.* (2006) |
| 4 | $\Delta lon \Delta marA$ | MG1655, *lon* : : *cat*, *marA* : : *kan* | Bhaskarla *et al.* (2016) |
| 5 | WT/VA | Isogenic complement strain for WT with plasmid pQE60, Amp<sup>r</sup> | This study |
| 6 | WT/p*marA* | Isogenic complement strain for WT expressing *marA* under T5 promoter present in plasmid pQE60, Amp<sup>r</sup> | This study |
| 7 | $\Delta marA$/VA | Isogenic complement strain for $\Delta marA$ with plasmid pQE60, Amp<sup>r</sup> | This study |
| 8 | $\Delta marA$/p*marA* | Isogenic complement strain for $\Delta marA$ expressing *marA* under T5 promoter present in plasmid pQE60, Amp<sup>r</sup> | This study |
| 9. | *E.coli* $\Delta lon::kan$ | Lon-deficient derivative of wild type | Kind gift from Dr. Nishad Matange, IISER (Pune, India) |
| 10 | *E.coli* $\Delta nfsA$ | *nfsA*-deficient derivative of wild type | Kind gift from Prof. Amit Singh, IISc (Bengaluru, India) |
| 11 | *E.coli* $\Delta nfsB$ | *nfsB*-deficient derivative of wild type | Kind gift from Prof. Amit Singh, IISc (Bengaluru, India) |

### Recording UV-Visible spectrum

To record the conversion products’ UV-visible spectra, the strains were grown in 5ml of MM media in the presence of 0.5Mm 2,4-DNP for 24 hr at 37°C at 160 rpm. The cells were pelleted down at 10,000 x g for 10 mins, and the supernatant was collected in a fresh falcon. The supernatant was filtered using a sterile syringe (BD, Belgium) with a PVDF filter membrane of 0.22 μm pore size (GE Health Care UK Limited, Hertfordshire, UK). Subsequently, 1:2 and 1:4 dilutions of the filtered supernatant were prepared by diluting with MM media. 0.5mM 2,4- DNP in MM filtered, post 24 hr of incubation following all the conditions of the bacterial cultures served as the reference. UV-visible spectra of the diluted supernatant solutions were recorded in a spectrophotometer (Spectramax M5e, Avantor, Pennsylvania, USA) using the software SoftMax pro-7. The spectra were plotted using the software Origin-2018.

### Quantification of UV-visible spectrum

The reduction in the concentration of 2,4-DNP during the conversion was quantified by calculating the drop in the absorbance of the peak maxima of 2,4-DNP, i.e. 364nm . The quantification was done using the formula (1), plotting the ratios in a graph.

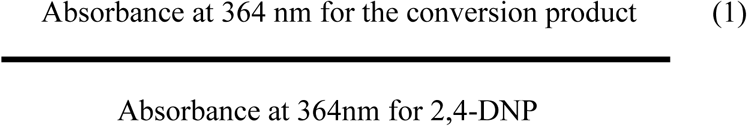

With more conversion product accumulation and reduction in the concentration of 2,4-DNP, the ratio will move more towards zero from one. i.e., one will be the ratio for DNP, which acts as the negative control in the experiment, and values will move towards zero with the reduction in the concentration of 2,4-DNP in the supernatant.

### LC-MS

The bacterial cultures were grown in 5ml MM with and without 2,4-DNP for 24 hr at 37°C at 160 rpm. The cells were pelleted down at 10,000 x g for 10 mins, and the supernatant was collected in a fresh falcon. The supernatant was filtered using a sterile syringe filter (Becton Dickinson) with a PVDF filter membrane of 0.22 μm pore size (GE Health Care UK Limited). LC-MS was performed in negative ionisation mode in LCESI Q Tof (Bruker Daltonics, Billerica, Massachusetts, USA). The compound spectra were analysed using Compass Bruker Data Analysis software, and fragments were analysed using the web tool Met-Frag. EGG was the database used during the fragment search.

### RNA purification, cDNA synthesis, and qRT-PCR

Total RNA was extracted from the WT and Δ*lon* cells grown for 3 hr, then treated with 0.5mM 2,4-DNP, and allowed to grow for 1 hr and 3 hr at 37°C and 160 rpm. Briefly, the cells were treated with TRIzol reagent (Sigma, St. Louis, Missouri, USA) for 1 hr at 37°C in a shaker dry bath, and the cells were lysed. The debris was pelleted down, the supernatant was transferred to fresh tubes and treated with chloroform (Sigma) and the organic phase was removed. The aqueous phase collected was treated with propanol (Sigma) to precipitate the RNA. The RNA was pelleted, washed with 70% chilled ethanol, dried and dissolved in 15 μL DEPC-treated water. RNA concentration and purity were quantified using a NanoDrop Spectrophotometer (Thermo Scientific, Waltham, MA, USA), followed by DNase treatment. Subsequently, the RNA was reverse transcribed to cDNA using Revert Aid (Thermo Scientific, Waltham, MA, USA). DNA contamination was tested by PCR amplification of *rrsC*. qPCR was carried out using the Bio-Rad CFX Connect System (Bio-Rad, CA, USA). The primers used are listed in Table S1. GapA was used as the reference gene for all conditions, and fold change was calculated using the 2^-ΔΔCt^ method. The primer efficiency for each primer pair was determined before performing PCRs, according to the MIQE guidelines (Livak and Schnittger 2001).

### Cloning of *marA* for trans-complementation

The calcium chloride method was used to prepare competent cells. Bacterial cultures were grown overnight from glycerol stock in the presence of antibiotics specific to the resistance cassette for the pre-inoculum. The pre-inoculum was normalised to O.D 2, and 0.2% of O.D 2 cells were cultured in 50ml flasks till the mid-log phase (O.D 0.4 – O.D 0.5). The flask was incubated at 4°C for one and a half hours.15 mL of the cell culture were pelleted down at 3000Xg for 15 mins at 4°C. The cells were resuspended in 7.5ml of filter sterilised 100mM MgCl_2_ (HiMedia), incubated at 4°C for 20 mins, and then pelleted down at 3000xg for 15 mins at 4°C. Following this, the cells were resuspended in 7.5ml of filter sterilised 100mM CaCl_2_ (HiMedia), incubated at 4°C for 20 mins, and then pelleted down at 3000Xg for 15 mins at 4°C. Then, the cells were gently resuspended in 1 ml of 100 mM CaCl_2_ in 15% glycerol.100μL aliquots were transferred to Eppendorf, flash-frozen with liquid nitrogen and stored at -80°C.

The WT MG1655 was used as the template for amplifying the *marA* gene using specific primers (Table S2) and Taq DNA polymerase (G-biosciences, Delhi, India). *marA* was targeted for cloning between EcoRI and HindIII sites in pQE60 plasmid using T4 DNA ligase (DX/DT, Bengaluru, India). The ligated pQE60 plasmid with the *marA* gene was amplified using the positive clones, which was confirmed by sequencing. The heat-shock method transformed the positive clones and control vectors into WT and Δ*marA*.

### Determination of Minimum Inhibitory Concentration (MIC)

MIC for the bacterial strains was performed using the broth dilution method. The bacterial cultures grown overnight at 37⁰C and 160rpm served as the pre-inoculum. 200μl of the LB was aliquoted in a 96-well microtiter plate (Tarsons, Kolkata, India), followed by the addition of different concentrations of ciprofloaxacin (HiMedia) ranging from 10 ng/ml to 100 ng/ml and tetracycline (HiMedia) ranging from 0.1μg/ml to 0.9 μg/ml. All the wells used 0.2% of OD2 culture as an inoculum. The plate was then incubated at 37℃ for 18hr followed by measurement of O.D. values at 600nm. The concentration of antibiotic is reduced to half is recorded as a the MIC value.

## Statistical Analysis

All statistical analyses were performed using ANOVA in GraphPad Prism 8, wherever applicable. Data were represented as mean ± S.D where * denotes p ≤ 0.05, ** denotes p ≤ 0.01, *** denotes p ≤ 0.001, and **** denotes p ≤ 0.0001.

## Acknowledgements

We thank the following: Preeti Bhatt, SR lab student, MRC, for helping with the spectral readings; Dr Nishad Matange, IISER Pune, for generously gifting the pBAD and p*lon* plasmids and WT and Δ*lon*: KR strains and Prof. Amit Singh for generously gifting Δ*nfsA* and Δ*nfsB strains.* We acknowledge the assistance for some experiments by Ms. Rithika Sri, St. Joseph University, Bengaluru. NSS is thankful to UGC for fellowship for graduate school studies. We are thankful for the DBT-IISc partnership grant and infrastructural support from the FIST program of the Department of Science and Technology, India. The authors of this study have no conflict of interest to declare.

## Notes

### Competing Interest Statement

The authors have declared no competing interest.

