## Supplemental Information for "Identification of 4-Amino-2-Nitrophenol as a novel inducer of phenotypic antibiotic resistance in *Escherichia coli*: roles of Lon protease and its substrate MarA"

**SI Table 1: List of primers used in the study**

| Sl.No | Primer name | Sequence (5'-3') |
|---|---|---|
| 1 | qRT *gapA* FP | TTTCCGTGCTGCTCAGAAAC |
| 2 | qRT *gapA* RP | GTCAACACCAACTTCGTCCC |
| 3 | qRT *nfsA* FP | GAACTTATTTGTGGCCATCG |
| 4 | qRT *nfsA* RP | TCACCAGTTCTTCACGTAAC |
| 5 | qRT *nfsB* FP | GGTTTATCTCAACGTCGGTA |
| 6 | qRT *nfsB* RP | GGTGTAGCCTTTCTCTTTCA |
| 7 | qRT*marA* FP | TGTCCAGGACGCAATACTGACG |
| 8 | qRT *marA* RP | TTTTGAAGGTTCGGGTCAGA |
| 9 | Comp *marA* FP | CCGGAATTCATGTCCAGACGCAATACTGACG |
| 10 | Comp *marA* RP | CCCAAGCTTGCGCGCCTAGCTGTTGTAATGATTTAAT |

**SI Table 2: List of plasmids used in the study**

| Sl No | Plasmid | Description | Reference |
|---|---|---|---|
| 1 | pQE60 | Low copy bacterial expression plasmid used for trans complementation from the constitutive T5 promoter. It contains an Ampicillin resistance cassette | Chandra *et al.*,2017;2020 |
| 2 | pBAD33 | Low copy number expression vector regulated by the arabinose operon | Matange,2020 |
| 3 | pBAD33-Lon | Plasmid for expression of Lon protease from an arabinose inducible promoter (Matange,2020) | Matange,2020 |

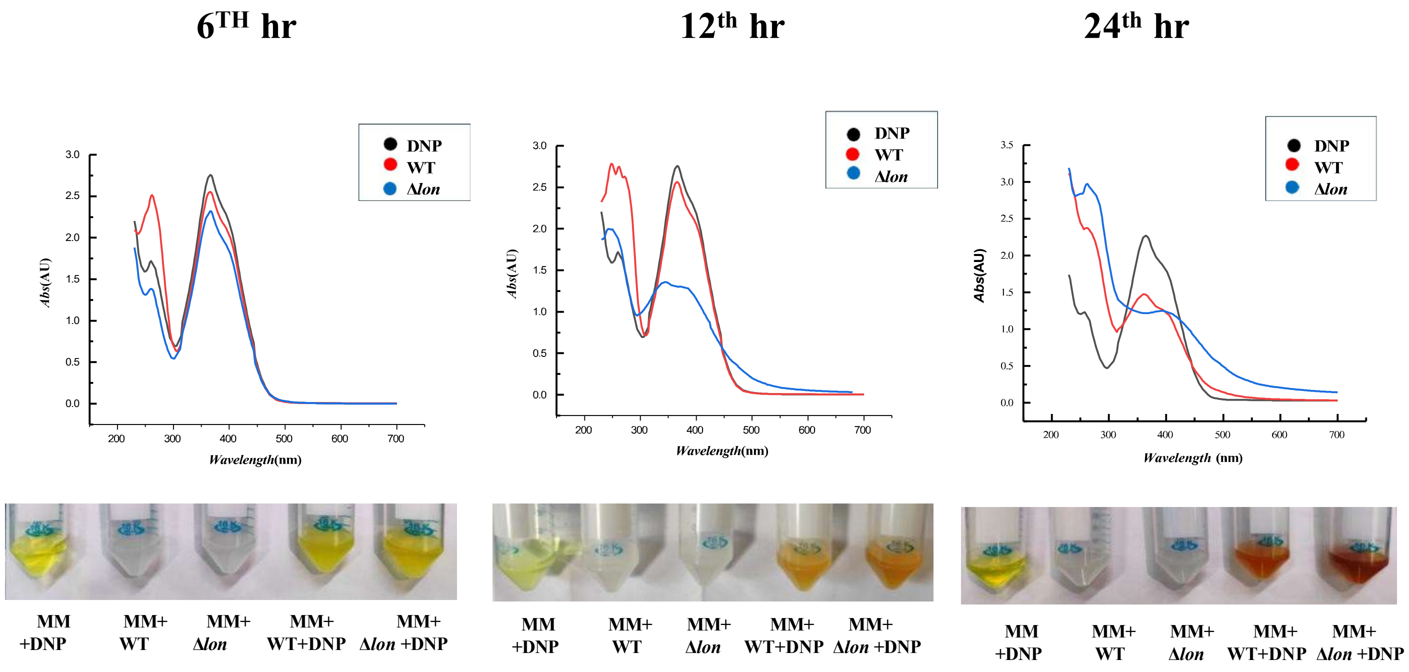


**SI Fig1. A reddish brown-coloured compound is produced in higher amounts in the Δ*lon* cells grown in minimal media in the presence of 2,4-DNP .***E. coli* MG1655 WT and Δ*lon* strains were cultured for 6,12, and 24 hrs at 37°C and 160 rpm in the presence of 0.5mM of 2, 4-DNP. Culture supernatants were collected from bacterial culture grown in the absence and presence of 0.5 mM 2,4-DNP. Subsequently, the UV-visible spectrum of the supernatants was measured after diluting with minimal media A) post 6 hr of treatment; B) post 12 hr of treatment; C) post 24 hr of treatment.

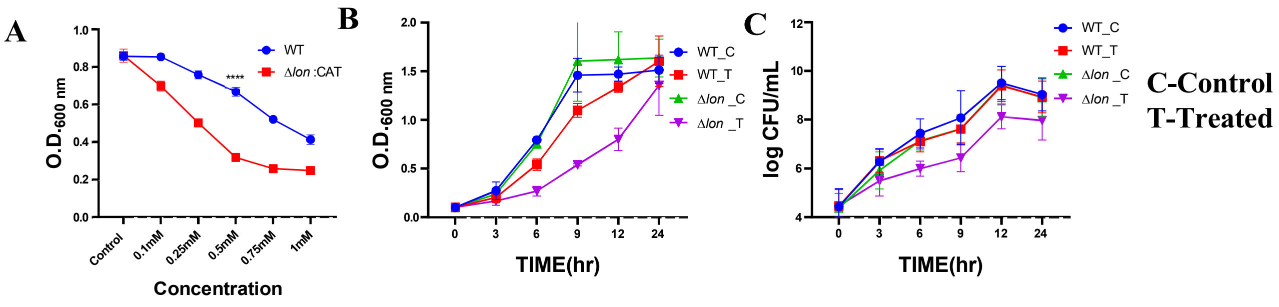


**SI Fig 2. The higher conversion product seen in the *lon* mutant is not due to higher growth in the presence of 2,4-DNP.** *E. coli* MG1655 WT and Δ*lon* strains were cultured for 6 hr at 37°C and 160 rpm in the presence of different concentrations of 2, 4-DNP. (A) Growth was assayed by measuring the O.D at 600 nm using a UV-visible spectrophotometer. B) Bacterial growth curve plotted as OD/ml v/s time and C) Bacterial growth curve plotted as log CFU/ml v/s time showing growth reduction in Δ*lon* strain in the presence of 0.5mM 2,4-DNP compared to WT. C represents the control group and T represents the treated group. The data are representative of three independent experiments. For statistical analysis, two-way ANOVA was performed, and the data were plotted as mean ± S.D where * indicates $p<0.05$. Statistical analysis was performed between WT and Δ*lon* strains for the different conditions.

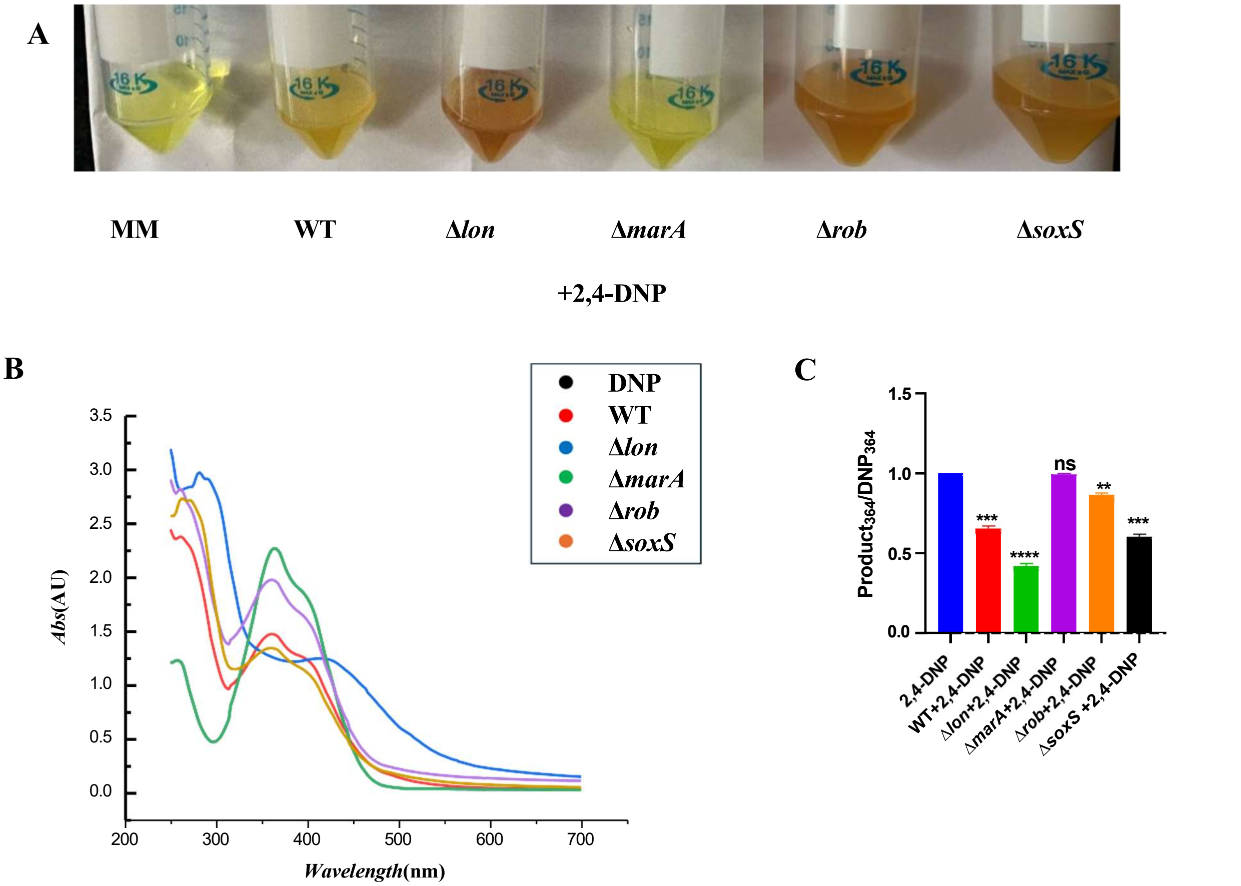


**SI Fig 3: *rob* and *soxS* mutants convert 2,4-DNP to the reddish brown-coloured compound.** *E. coli* WT, Δ*lon*, Δ*rob* and Δ*soxS* were cultured in the presence of 0.5mM 2, 4-DNP for a period of 18 hr at 37$^0$C and 160 rpm. (A)Tubes showing conversion product formed post 18hr of treatment;(B) UV-visible spectrum of the supernatant after diluting 1:2; (C) graph showing quantification of spectrum. The data are representative of three independent experiments plotted as mean ± SD. * indicates P<0.05. Statistical analysis was performed for each strain relative to its untreated control. Comparison between the strains is indicated wherever significant.

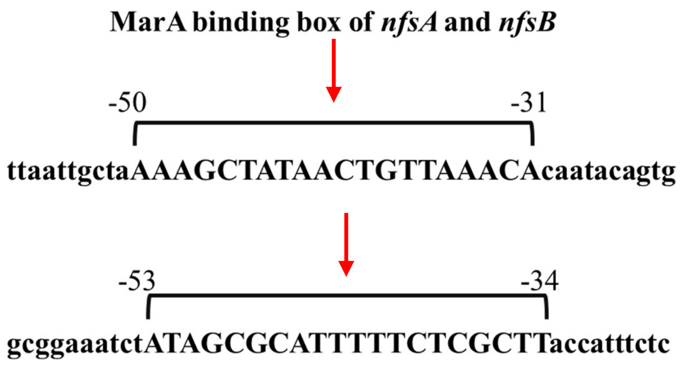


**SI Fig 4 :Representative image of Mar A binding box of *nfsA* and *nfsB* :**Position of Site Center Relative to Transcription Start Site for *nfsA* and *nfsB* (bp) are -40.5 and -43.5 respectively (marked with red arrow)
